# Extracellular domain, hinge, and transmembrane determinants affecting surface CD4 expression of a novel anti-HIV chimeric antigen receptor (CAR) construct

**DOI:** 10.1101/2023.10.25.563930

**Authors:** Giorgio Zenere, Chengxiang Wu, Cecily C. Midkiff, Nathan M. Johnson, Christopher P. Grice, William C. Wimley, Amitinder Kaur, Stephen E. Braun

## Abstract

Chimeric antigen receptor (CAR)-T cells have demonstrated clinical potential, but current receptors still need improvements to be successful against chronic HIV infection. In this study, we address some requirements of CAR motifs for strong surface expression of a novel anti-HIV CAR by evaluating important elements in the extracellular, hinge, and transmembrane (TM) domains. When combining a truncated CD4 extracellular domain and CD8α hinge/TM, the novel CAR did not express extracellularly but was detectable intracellularly. By shortening the CD8α hinge, CD4-CAR surface expression was partially recovered and addition of the LYC motif at the end of the CD8α TM fully recovered both intracellular and extracellular CAR expression. Mutation of LYC to TTA or TTC showed severe abrogation of CAR expression by flow cytometry and confocal microscopy. Additionally, we determined that CD4-CAR surface expression could be maximized by the removal of FQKAS motif at the junction of the extracellular domain and the hinge region. CD4-CAR surface expression also resulted in cytotoxic CAR T cell killing of HIV Env^+^ target cells. In this study, we identified elements that are crucial for optimal CAR surface expression, highlighting the need for structural analysis studies to establish fundamental guidelines of CAR designs.

## Introduction

Chimeric antigen receptors (CAR) are recombinant fusion proteins designed to mimic T cell receptor signaling and redirect immune functions against desired antigens [1, 2]. CARs have the advantage of bypassing canonical MHC presentation and restrictions on T cell stimulation [3, 4]. The structure of a CAR has four major components: an extracellular antigen recognition domain(s), a hinge region, a transmembrane domain to anchor the receptor to the cell surface, and intracellular signaling domains to drive cell activation and confer immune function. T cells derived from patient blood and engineered with CARs have been used to successfully target tumor antigens, as seen by the high reduction in remission rates reported against hematological cancers such as acute lymphoblastic leukemia and non-Hodgkin lymphomas [5–12]. However, despite the progress that has been made in treating hematological malignancies, many challenges remain for successful CAR T cell therapy of solid tumors and chronic HIV infection [13, 14].

To improve the efficacy of CAR T cells in these fields, novel CAR structures are being designed and evaluated. These often include the generation of new extracellular domains [15–18], hinge regions taken from different receptors [19], swapping transmembrane domains or intracellular domains [20, 21], and even arming CARs with cytokine receptors or knocking out PD-1 expression [22–24]. In the instance of HIV immunotherapy, anti-HIV CAR T cells were first designed with the full-length whole CD4 extracellular domain linked to an intracellular TCRζ chain [16, 25–32]. The full-length CD4 extracellular domain was originally chosen because of its inherent advantage of recognizing the primary receptor-binding site on HIV envelope glycoproteins, which must be retained on all clinical HIV-1 variants [33–36]. Since then, researchers have developed a truncated version of CD4 containing only immunoglobulin domain 1 and 2 (D1D2) to improve CAR safety while retaining CD4 binding affinity to HIV envelope glycoprotein [37]. This is achieved because HIV envelope glycoprotein binding to CD4 only requires D1D2 whereas the immunoglobulin domains 3 and 4, lacking in the novel CD4 CAR extracellular domain, are involved in CD4 oligomerization for stable binding to the MHC class II molecule [38–40]. Therefore, D1D2 CD4 CAR have reduced potential off-target effects compared to whole CD4 CARs. Furthermore, other anti-HIV CARs demonstrated that swapping CD4-CAR transmembrane domain (TM) for CD8α TM domain decreased CAR homology to the HIV cellular receptor and reduced the susceptibility of CD4- expressing CAR T cell to HIV infection [41].

Although some of the latest strategies look promising, one potential reason why anti-HIV CARs have yet to show clinical benefits is because some of the domains incorporated in the new CAR structures have motifs whose biochemical function and structural importance are still poorly understood and can dramatically affect the success of a CAR strategy. In the present study, we first attempted to generate a novel CD4 based CAR that combined the truncated CD4 D1D2 extracellular domain with the innovative CD8α TM to improve the safety and efficiency of anti-HIV CAR. We observed a lack of surface detection by flow cytometry and confocal microscopy but determined that the CAR was synthesized and detectable intracellularly. Through a series of rescue vectors, this study identified specific CAR elements that are crucial for optimal CAR surface expression and maintained cytolytic activity. Our findings illustrate the need for thorough analysis of the CARs structure to help establish fundamental guidelines of CAR designs that will help the field generate effective therapies.

## Results

### Newly designed D.66.α CD4-CAR is not detectable on the cell surface despite being synthesized

In an attempt to combine two CAR structural domains within the same chimeric protein, we generated a retroviral vector (D.66.α) using the truncated CD4 extracellular domain containing immunoglobulin domains 1 and 2 (D), based on the GenBank database [42], linked with a 66 amino acid hinge (D.66) and TM domain from CD8α (α) [37, 41] (Figure 1). Our goal was to compare retroviral expression of D.66.α and C.39.28, a full-length CD4 extracellular domain (C) with a 39 amino acid hinge (39) and TM domain from CD28 (28) that has known surface expression and functional capabilities [26, 28, 43]. Retroviral transduction of HEK 293T cells produced various clones with integrated CAR vector (Supplemental Figure 1a). However, CD4 surface expression of the newly generated D.66.α CAR was not detectable by flow cytometry. In contrast, CD4 surface expression was robust with the C.39.28 CAR (Figure 2a). To eliminate the possibility that the vector failed to express because of a possible defect in vector integration or viral particle production, HEK 293T cells were transfected by calcium phosphate method with the same vectors. Similar to the transduction experiment, cells transfected with C.39.28 expressed CD4 CAR on the surface while cells transfected with D.66.α did not (Figure 2b). The lack of surface detection in the D.66.α CAR was verified by staining with an anti-Myc antibody (Supplemental Figure 1b and 1c). Furthermore, proviral vector DNA was detectable in individual transduced clones by qPCR; however, these clones did not have detectable CAR surface expression (Supplemental Figure 1a).

**Figure 1.**
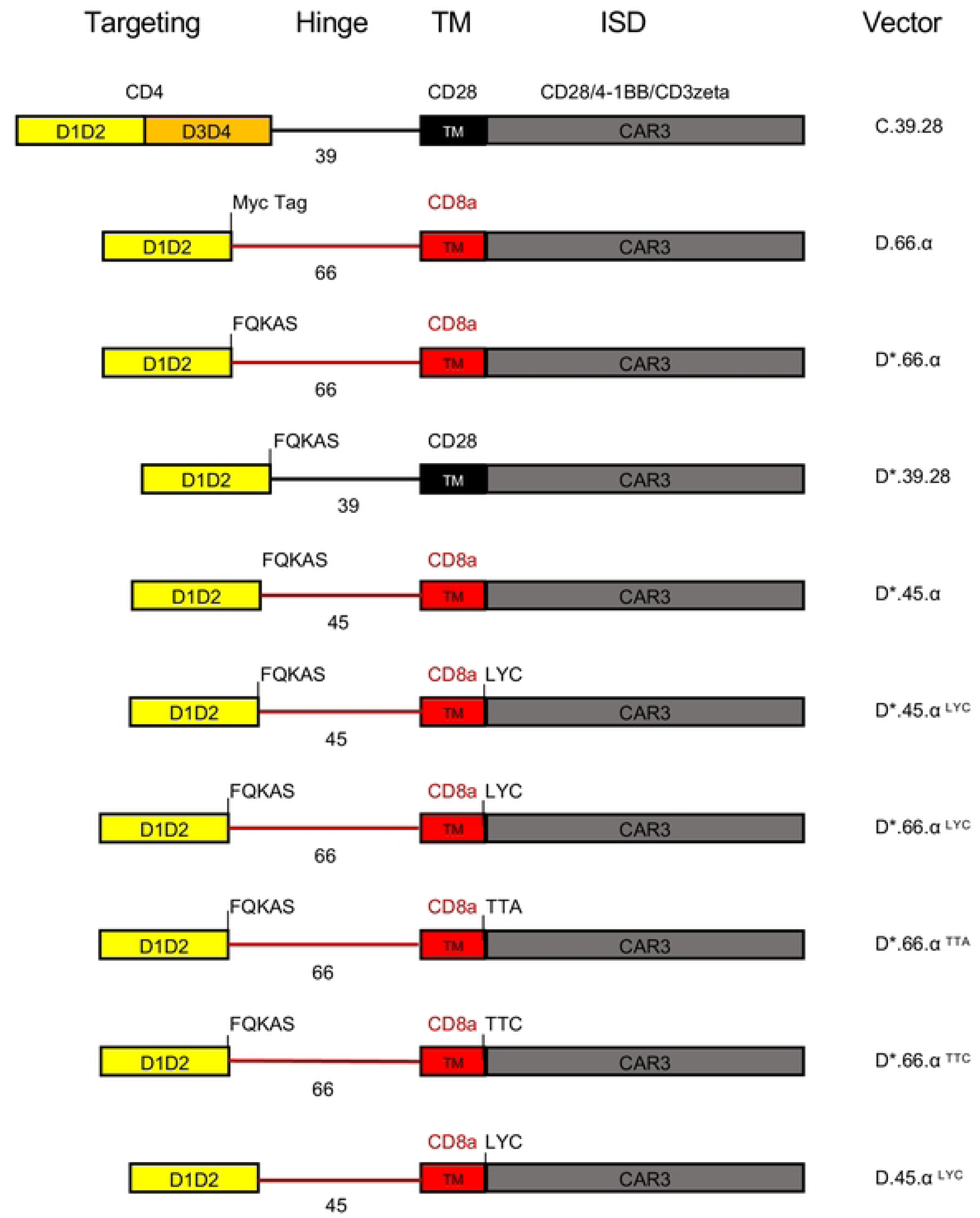
Schematics representing the CAR vector maps for each of the major constructs used in this study. CD4 extracellular domain is designated as domain 1 and 2 (yellow), and domain 3 and 4 (orange). Black hinge and TM indicate a CD28 origin; red hinge and TM indicate CD8α origin. Intracellular domains are indicated at the top. Vectors were named based on extracellular configuration, length of hinge, transmembrane domain used, and motifs included.

**Figure 2.**
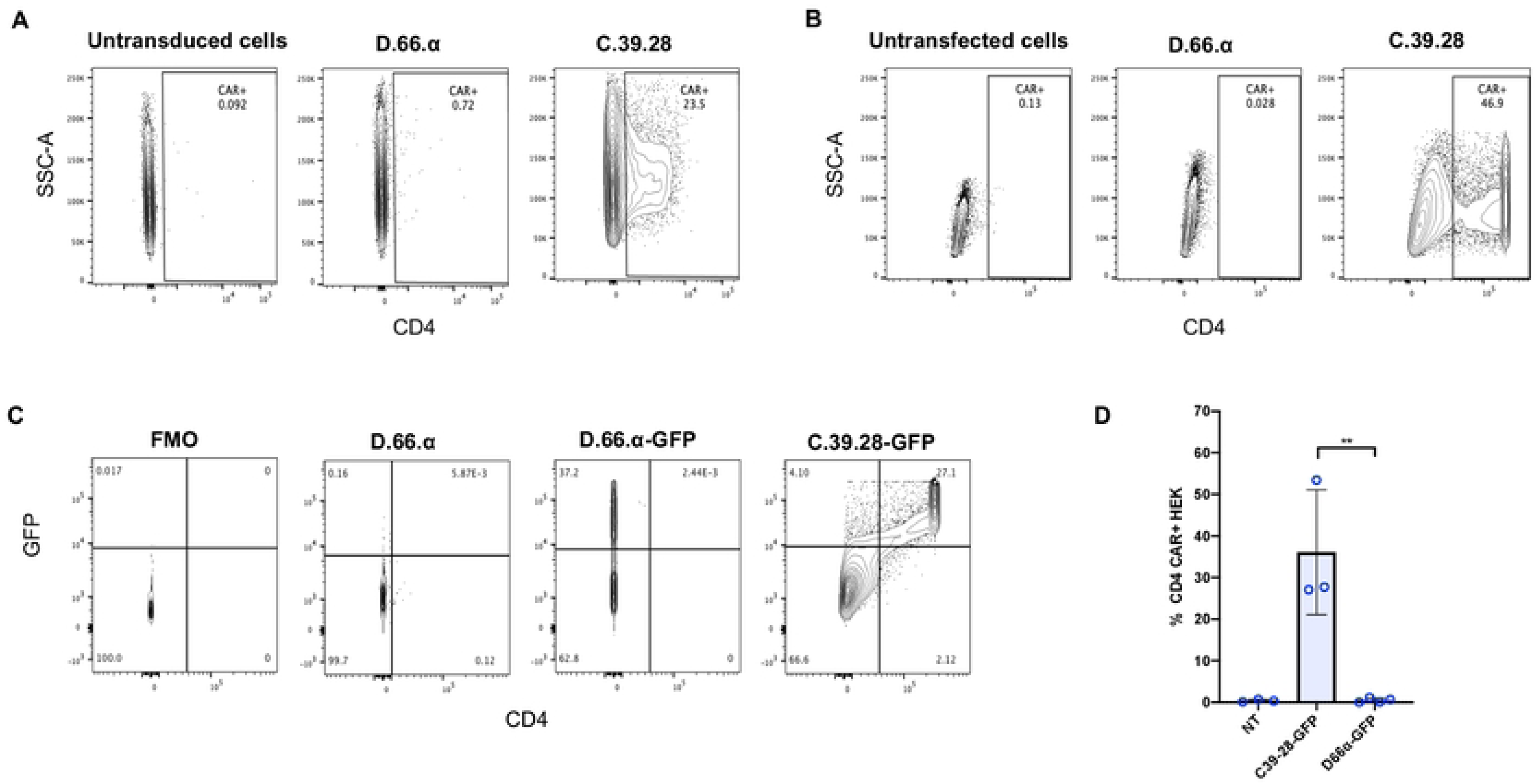
Novel D1D2 CAR is not expressed on the cell surface but the protein is synthesized. **A.** Representative flow cytometry plots of CD4 CAR expression on the surface of vector-transduced HEK293 T cells. In contrast to the C.39.28 CAR vector, no surface expression was detected on cells transduced with D.66.α. **B.** Representative flow cytometry plots of CD4 CAR expression on the surface of HEK293 T cells transfected by calcium-phosphate method. No surface expression was detected on cells transfected with D.66.α. **C.** Representative flow cytometry plots of CD4 CAR expression on the surface of vector-transduced HEK293 T cells (x axis) and GFP expression from the second gene (y axis). Dual CD4 CAR and GFP expression was observed with positive control C.39.28-GFP but not with D.66.α-GFP. **D.** Histogram representing the frequency of transfected HEK 293T cells expressing surface CD4 CAR-GFP. Statistical analysis was performed as unpaired parametric two-sample t test; a significant difference in intracellular detection was observed between groups (** = p ≤ 0.01).

Since the newly designed D.66.α CAR was integrated in the host genome of transduced cells but was not detected on the surface by either transduction or transfection, we hypothesized that the CAR protein was being synthesized but not express on the cell surface. To demonstrate that CAR was being synthesized, we generated bicistronic vectors to express a second gene encoding a green fluorescence protein (GFP) (Supplemental 1D). GFP was chosen as a reporter gene because its detection via flow cytometry does not require surface expression but only its translation. After transfection with D.66.α-GFP, GFP expression was detected in cells but CAR surface expression was still not detected. In contrast, cells transfected with C.39.28-GFP expressed both CD4 CAR and GFP (Figure 2c and 2d). Since the bicistronic D.66.α-GFP vector included a TPT2a domain to ensure that both proteins were synthesized *en block* [44], these results suggested that the D.66.α CAR was being synthesized but not expressed on the cell surface.

### D1D2-66H-8 αTM CAR is expressed intracellularly but not on the cell surface

Once we established that the D.66.α CAR protein was being synthesized but still lacked surface expression, we set out to determine if we could detect intracellular expression of the CAR with the same antibody that showed robust surface detection of the C.39.28 CAR. Therefore, we adopted our intracellular staining protocol to detect CD4 CARs and observed intracellular expression of the D.66.α CAR in transfected HEK 293T cells (Figure 3a). However, there was significantly less intracellular expression of D.66.α than C.39.28 (Figure 3a and 3b). This difference in intracellular and extracellular expression was also observed when overall data for monocistronic and bicistronic CARs were combined (Figure 3c and 3d). Therefore, these results confirmed that D.66.α lacked surface expression but was detectable intracellularly.

**Figure 3.**
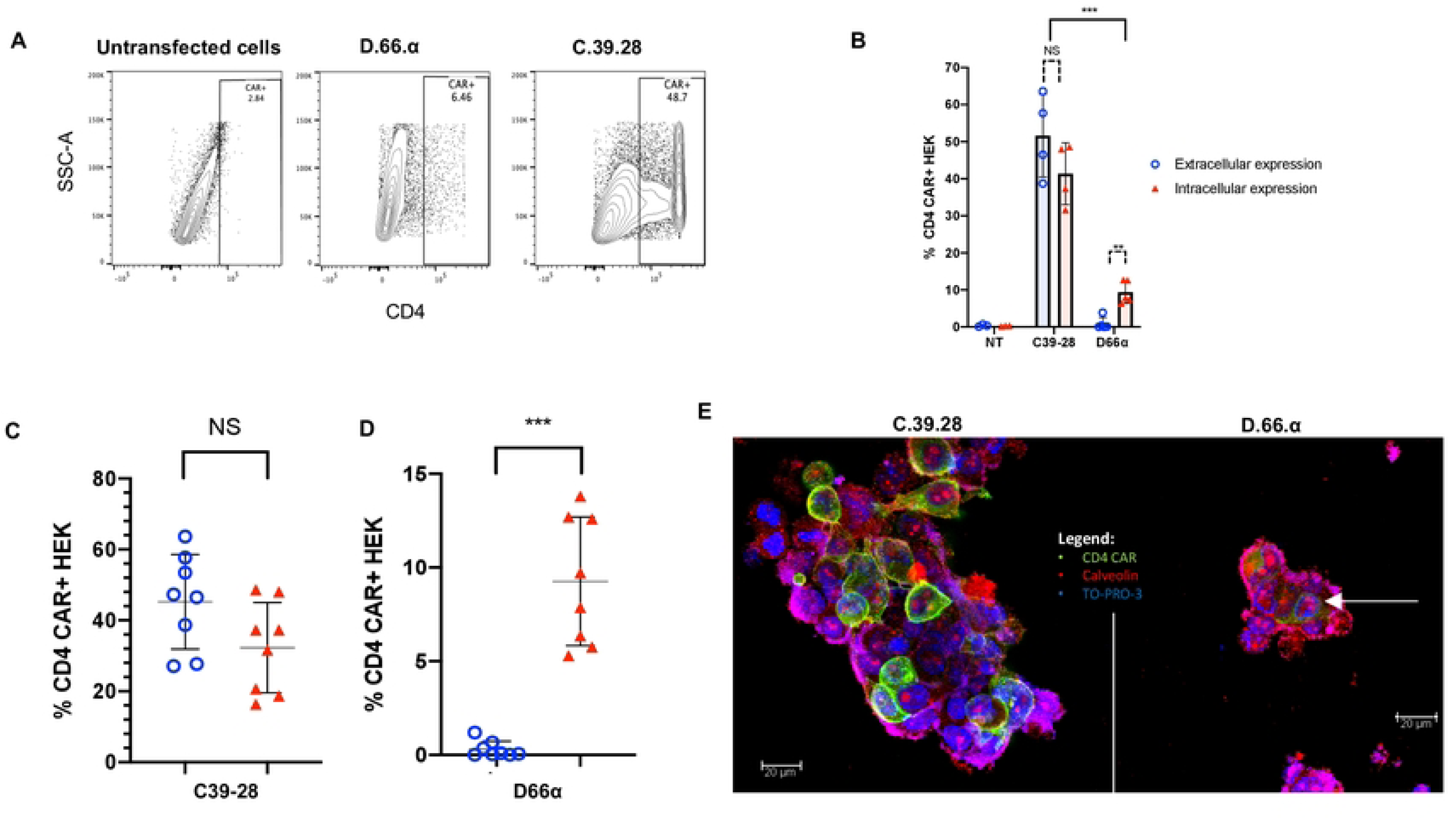
D1D2 CAR is detected intracellularly but not on cell surface. **A.** Representative flow cytometry plots of CD4 CAR intracellular expression in transfected HEK293 T cells. Reduced but significant intracellular expression of D.66.α was detected compared to C.39.28 vector. **B.** Histogram representing the frequency of transfected HEK 293T cells expressing surface CD4 CAR (red filled triangle) and intracellular CD4 CAR (blue circle) expression. Significant difference in CAR detection was observed between the vectors (p<0.0001). No significant difference in surface and intracellular-expressing cells was detected for the positive control C.39.28 (p= 0.191). Significant difference between surface and intracellular expression was observed in the D.66.α CAR (p= 0.0066). **C&D.** Scatter plot for CD4-CAR expression in transfected HEK 293T cells when monocistronic and bicistronic vectors are grouped. Robust detection of C.39.28 was observed on the surface and intracellularly (p=0.098). Significant, but comparatively reduced detection of intracellular D.66.α expression was observed with no surface expression (p=0.0002). **E.** Images of CAR-transfected HEK 293T cells taken by confocal microscopy. Blue is TO-PRO-3 representing nuclear stain; green is CD4-CAR; and red is caveolin representing the plasma membrane. C.39.28 CAR staining shows green expression above the red stain and around the cell, suggesting surface expression. D.66.α CAR staining shows green expression surrounded by red, suggesting intracellular expression only.

Finally, to confirm the CAR expression results observed by flow cytometry, HEK 293T cells transfected with either D.66.α or C.39.28 CAR were evaluated by confocal microscopy using the same anti-CD4 antibody. Confocal microscopy images showed that C-39-28 CD4-CAR (green) was detectable around and beyond the caveolin-stained plasma membrane (red), whereas the D.66.α CD4-CAR stained below the cell membrane (Figure 3e). Consequently, these results complement our findings by flow cytometry.

### Extracellular CD4 domain is not involved in inhibiting new CAR surface expression

Once it was confirmed that the new D.66.α CD4-CAR was synthesized and detectable intracellularly but without stable cell surface expression (Figure 3), we investigated the structural reasons behind the lack of CAR surface expression. Compared to C.39.28, which strongly expressed extracellularly and intracellularly, the D.66.α CAR had three major modifications; (i) the full-length CD4 extracellular domain was truncated to immunoglobulin domains 1 and 2, (ii) the length of the hinge, and (iii) TM domains were changed to CD8α hinge and TM domain respectively. As a result, D.66.α had a longer hinge (66 amino acids compared to the 39 amino acids present in the CD28 hinge). In addition, there were several other modifications that were added (including a Myc tag), but those were shown to have no impact on recovery of CAR surface expression (Supplemental Figure 2a and 2b).

To determine if the defect causing the lack of CAR surface expression was found in the truncated extracellular domain, we compared the D1D2 sequence [42] with other D1D2 sequences used in other CD4-CARs and soluble CD4 inhibitors (1-183 aa) [37, 45]. Our CAR terminated at the end of immunoglobulin domain 1 and 2 [42], while other D1D2 CAR domains (D*) incorporated the first 5 amino acids (FQKAS) of domain 3 [37] (Figure 1). Nevertheless, addition of the FQKAS motif to our non-expressing D.66.α CAR (D*.66.α) did not recover surface expression (Supplemental Figure 2c and 2d). In fact, there were no differences in extracellular or intracellular expression between D.66.α CARs with or without FQKAS added to the end of the D1D2 (Supplemental Figure 2c).

To further understand if the extracellular domain was inhibiting CAR surface expression, the entire hinge and CD8α transmembrane domain of the D*.66.α CAR was replaced with a shorter CD28 hinge and a CD28 TM (D*.39.28; Figure 1). Results showed that when the CD28 hinge and TM domain were included, the D*.39.28 recovered CAR surface expression compared to D*.66.α (p=0.01) (Supplemental Figure 2c) suggesting that the D1D2 CAR with a shorter hinge (39 base pair of CD28 instead of 66 bp of CD8α) and CD28TM recovered CAR surface expression compared to its predecessor (p=0.01) (Supplemental Figure 2c). Additionally, overall CD4 expression with the D*39.28 CAR was still reduced compared to C.39.28. These results were confirmed by confocal microscopy, where expression of the D*39.28 (green) is visible on and around the plasma membrane of cells (red) while expression of the D*.66.α CAR is only visible within the cell membrane (Supplemental Figure 2d). These data highlighted that the major issue preventing CAR surface expression was found in the hinge and transmembrane domain region and not in the truncated CD4 extracellular domain.

### Hinge length partially affects CAR surface expression

To test if the hinge length was responsible for the lack of CAR surface expression, we shortened the hinge of the non-surface expressing D*.66.α CAR from 66 amino acids (aa) to 45 aa (D*.45.α Figure 1), as previously published [41]. Interestingly, shortening of the CD8α hinge from 66 to 45 aa resulted in partial recovery of CAR surface expression while maintaining the same level of intracellular detection (Figure 4a). Although surface CD4 detection of D*.45.α CAR was robustly increased compared to D*.66.α, it was deemed partial because it was still significantly lower than D*39.28 CAR (Figure 4a and Supplemental Figure 2c). These results suggested that a longer hinge length contributes to the inhibition of CAR surface expression.

**Figure 4.**
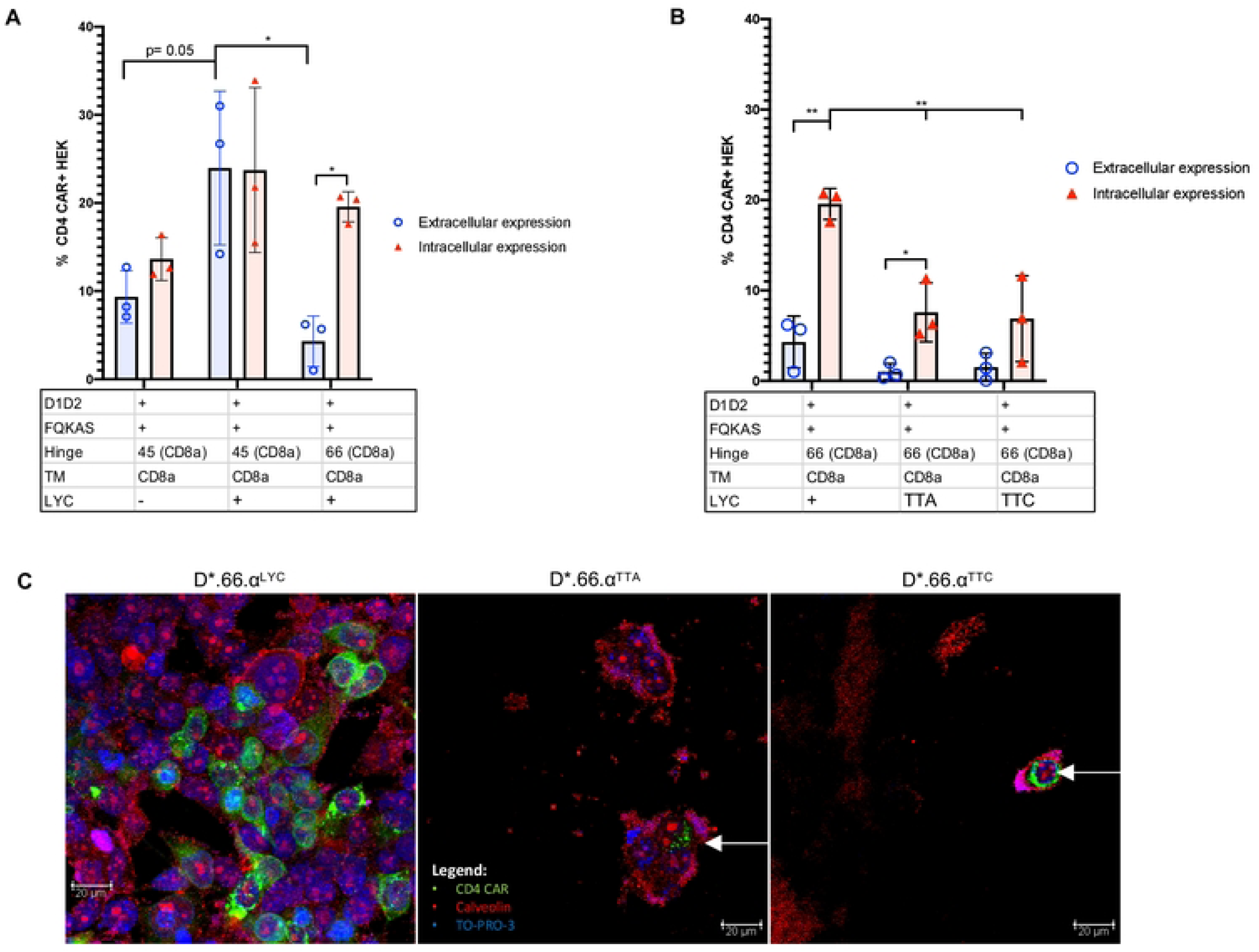
Shortening of the CD8α hinge length and LYC motif are required for D1D2 CAR surface expression. **A.** Histogram representing the frequency of transfected HEK 293T cells expressing surface CD4 CAR and intracellular CD4 CAR for the various vectors. Positive CAR detection was observed on the surface of cells transfected with a shorter hinged CAR (p= 0.01); however, expression was still reduced compared to control CAR (p= 0.002). Intracellularly, there was no change in CAR detection between long and short hinge (p= 0.227). Enhance frequency of CAR surface detection was observed on cells transfected with CAR that included the LYC motif (p= 0.05); however, expression was significantly reduced if the construct containing a LYC motif also included a longer CD8α hinge (p=0.02). **B.** Histogram representing the frequency of HEK 293T transfected with CARs containing the longer CD8α hinge to eliminate any variability/benefit given by a shorter CD8α hinge. When LYC motif was mutated to TTA or TTC, total CAR expression was significantly reduced (p= 0.01 and p= 0.03) and more intracellular CAR expression than surface expression (p=0.04). **C.** Images of CAR-transfected HEK 293T cells taken by confocal microscopy. Blue represents TO-PRO-3 nuclear staining; green represents CD4-CAR staining; and red represents caveolin plasma membrane staining. Differences in confocal surface CAR detection were observed, with D*.66.α^LYC^ expressed on the cell surface but D*.66.α^TTA^ and D*.66.α^TTC^ confined intracellularly. Statistical analyses were done using unpaired parametric two-sample t-test.

### Both the presence of the LYC motif at the end of CD8αTM and shortened hinge are responsible for recovering CAR surface expression

Since shortening of the hinge only partially, but distinctively, recovered CAR surface expression, we hypothesized that there must be a second defect, likely found in the CD8α transmembrane domain (TM), that prevents robust CAR surface detection. Proteomics analysis of the CD8α TM sequence in the D*.45.α CAR, compared to the CD8α TM sequence found in other published CARs [41], revealed a difference at the end of the TM; specifically, it included LYC, the first 3 amino acids of the CD8α intracellular domain in the CD8αTM reference sequence used in our CARs [46]. Addition of the missing LYC motif at the end of CD8αTM (D*.45.α^LYC^) strongly recovered CAR surface expression to levels similar to D*.39.28 (Figure 4a and Supplemental Figure 2c). Therefore, these results demonstrated that the addition of LYC motif from the beginning of the intracellular domain was crucial for robust CAR surface expression.

To determine if the LYC motif alone was sufficient to recover CAR surface expression, the 45 aa hinge on D*.45.α^LYC^ (which expressed on the surface) was replaced with the 66 aa hinge found on non-surface expressing CARs (Figure 1 and Figure 4a). Transfection with D*.66.α^LYC^ resulted in significant reduction of CAR surface expression but intracellular expression was comparable to D*.45.α^LYC^ (Figure 4a). These results demonstrated the importance of CD8α hinge length for CAR surface expression and showed that both a shorter CD8α hinge and the addition of LYC motif at the end of CD8α TM act synergistically to increase surface CD4-CAR expression.

To further understand the importance of the LYC motif, we took the D*.66.α^LYC^, which had strong intracellular expression but reduced extracellular expression, and mutated the LYC motif to TTA or TTC in an attempt to contrast the possible biochemical interactions of the residues (Figure 1 and 4b). Interestingly, mutating the LYC motif at the end of CD8α TM to TTA and TTC abrogated CAR surface expression, but also significantly reduced intracellular CAR expression (Figure 4b). Therefore, these experiments further highlighted the importance of the LYC motif for proper CAR detection. Results were also confirmed by confocal microscopy, where mutating the LYC motif distinctively resulted in a complete loss of CAR surface expression, as CAR expression with the TTA and TTC variants were confined to the intracellular compartment (Figure 4).

### The extracellular FQKAS motif reduces CAR detection

To investigate why the D1D2 CD4-CAR variations that strongly recovered surface expression (mainly D*.39.28 and D*.45.α^LYC^) were still not as efficiently and broadly expressed as the full-length extracellular domain CD4 construct C.39.28, we removed the FQKAS domain from D*.45.α^LYC^ to produce a D.45.α^LYC^ CAR (Figure 5). Interestingly, removal of the FQKAS domain at the end of the extracellular domain 2 recovered the same level of efficient CAR expression as C.39.28 in transfected HEK 293T cells (Figures 5 and 6). Therefore, our results suggest that the presence of FQKAS reduced overall CAR expression, both extracellularly and intracellularly.

**Figure 5.**
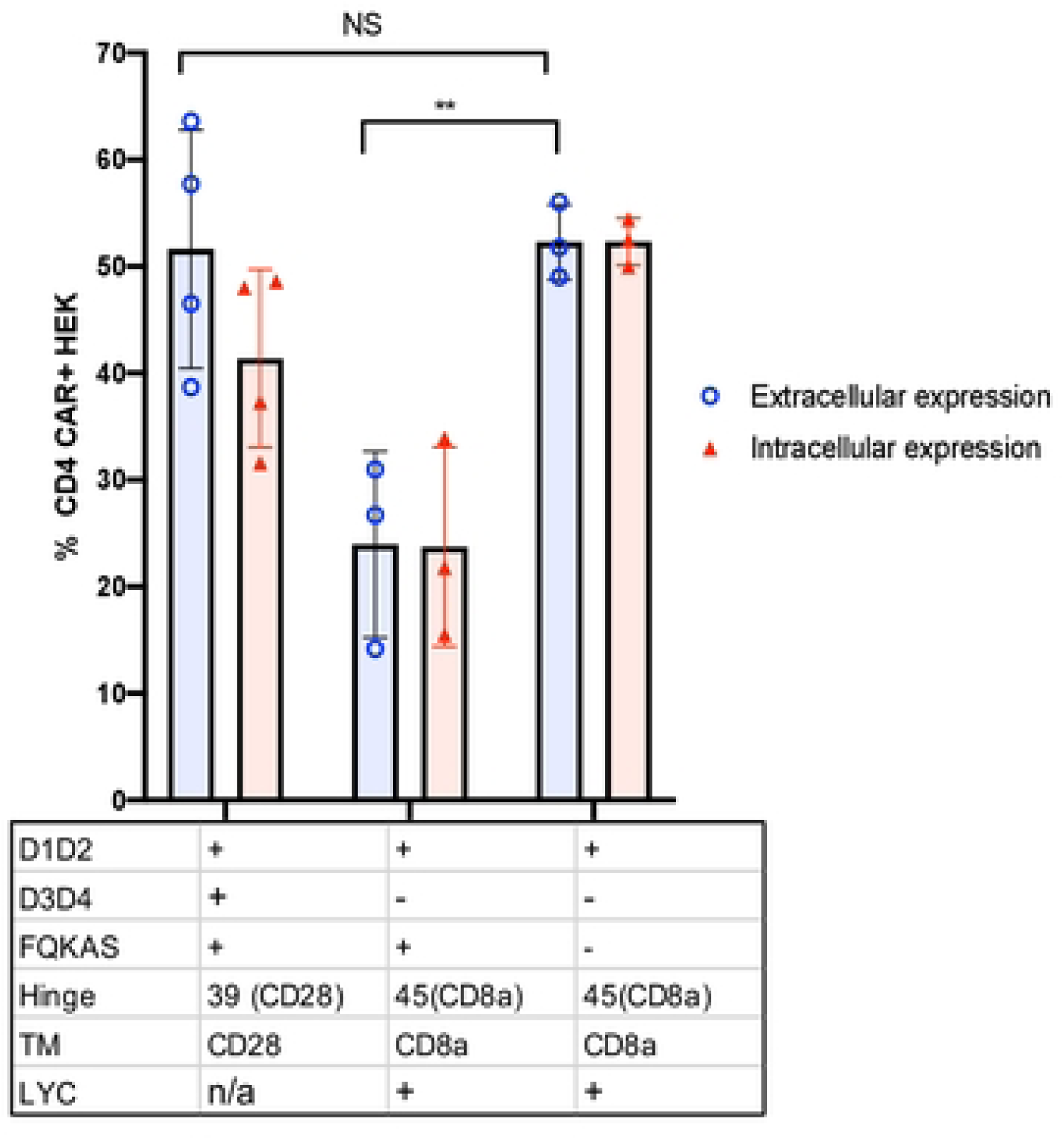
FQKAS motif hinders but does not inhibit CAR surface expression. Histogram representing the proportion of transfected HEK 293T cells expressing surface CD4 CAR and intracellular CD4 CAR. Positive CAR expression was observed on the surface of all transfected cells, but the frequency was significantly higher in vectors that did not incorporate FQKAS (p= 0.006). Statistical analysis was done using unpaired parametric two-sample t-test.

**Figure 6.**
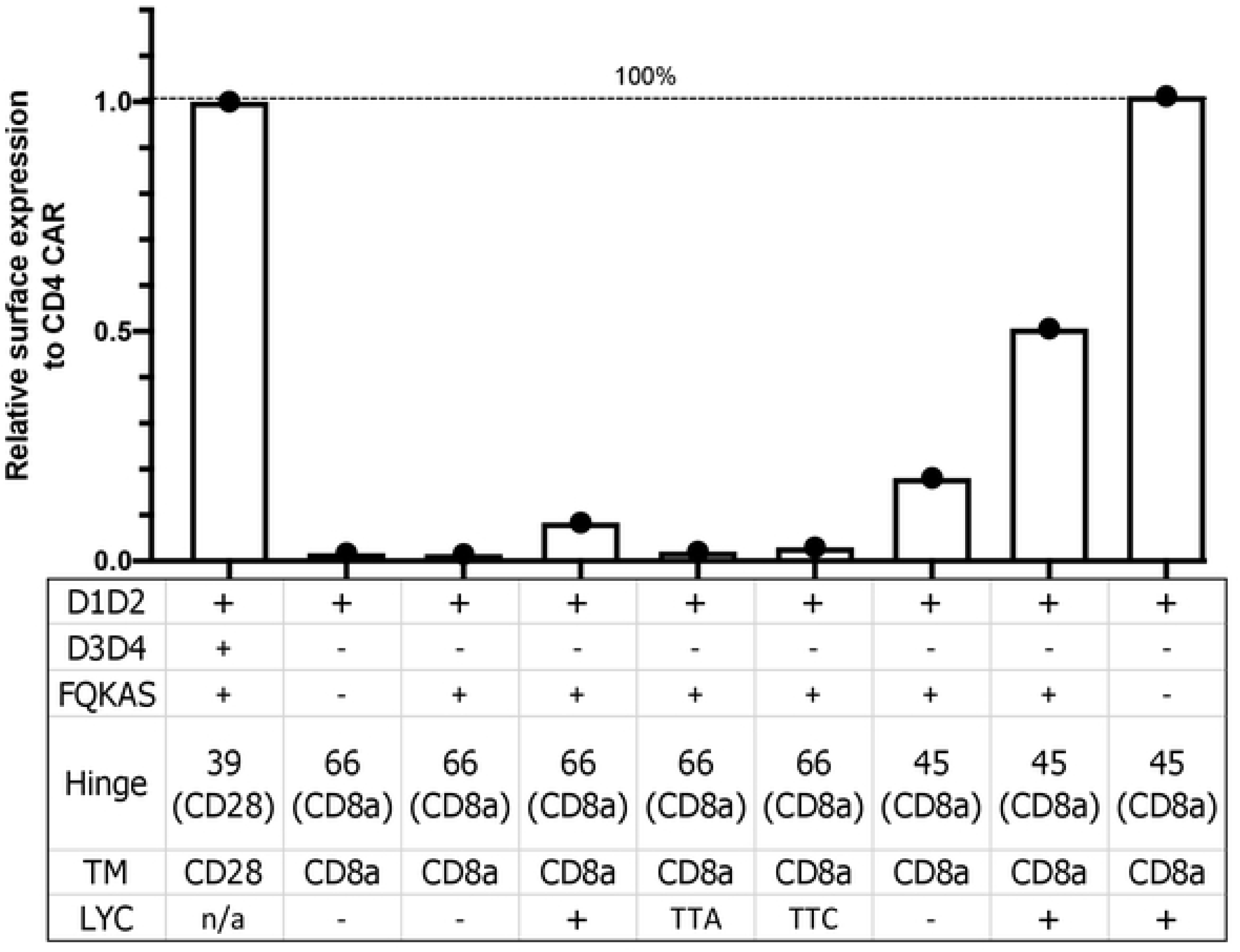
Summary graphic of variant CAR surface expression. Data analysis was based on the relative surface expression of positive control, whole CD4 CAR. Addition of LYC motif and shortening CD8α hinge each partially recovered CAR surface expression, which was enhanced when both modifications were combined. Removal of motif FQKAS improved CAR surface expression to maximal levels seen in the positive control.

### Recovered CAR expression enhances cytotoxic CAR T cell activity

Once we recovered surface expression of D.45.α to level similar to C.39.28, we set out to determine that the CAR had similar functional abilities as well. Since cytotoxic killing is one of the main read out of CAR T cell ability, we compared cytotoxic ability of D.45.α to C.39.28 CAR T by co-incubating cells with HIV+RFP+ U1 target cells (Figure 7). While untransduced T cells demonstrated limited killing ability at various E:T ratios, D.45.α CAR T cell exhibited similar levels of target killing as C.39.28 CAR T cells (Figure 7). Therefore, our data demonstrates that the 2 CAR vectors with similar surface expression also maintained CTL activity, the primary CAR T cell functional activity.

**Figure 7.**
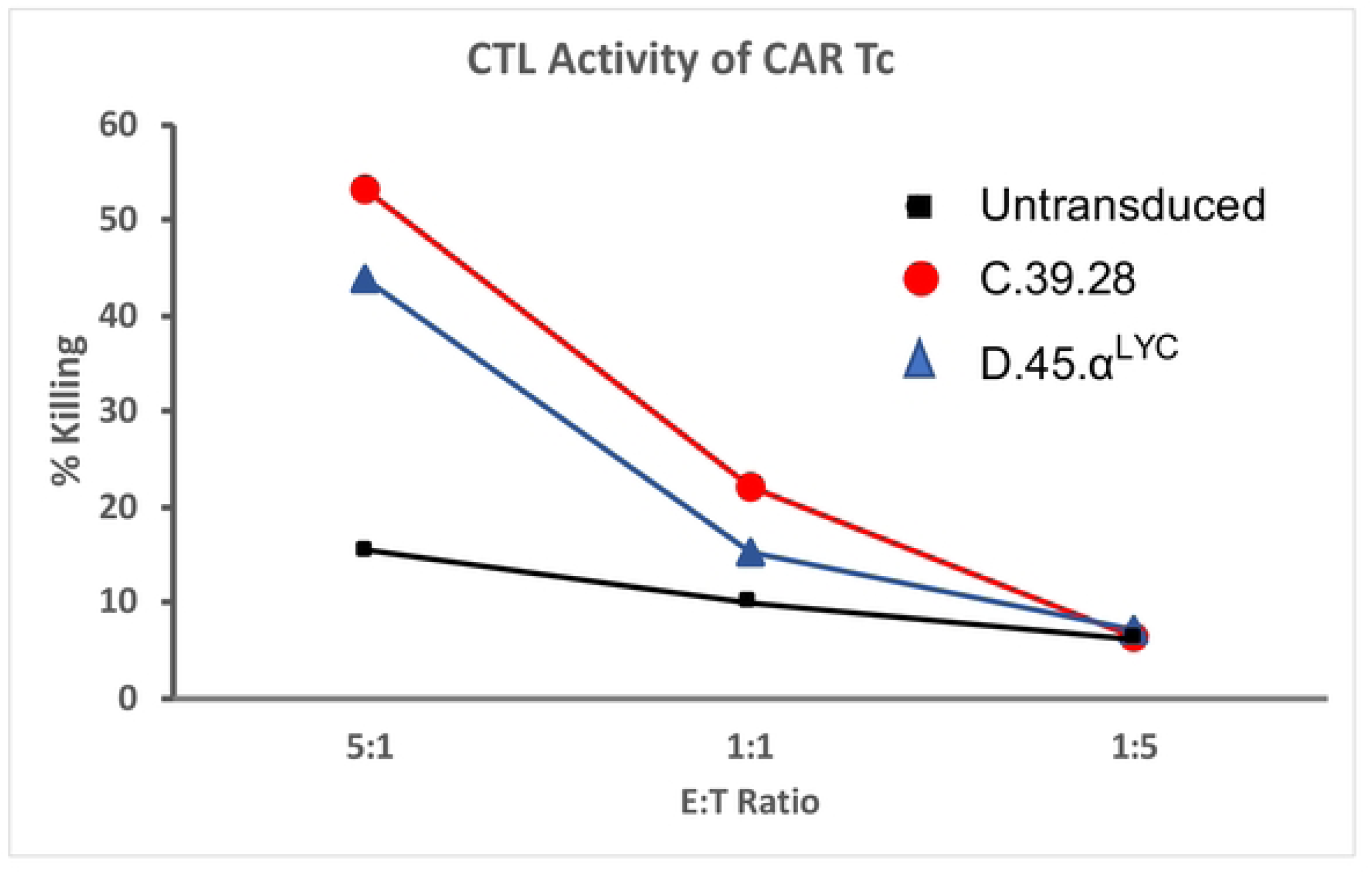
Cytolytic activity of CAR T cells. Primary human T cells were transduced with select vectors and incubated for 24 hours with HIV+ target cells at different E:T ratios. Untransduced T cells showed limited cytotoxic abilities whereas T cell transduced with D.45.α^Lyc^ showed similar killing capacity as the full length CD4 CAR C.39.28

## Discussion

Overall, our study started by evaluating a novel CD4-CAR design that lacked surface expression but showed intracellular expression, as determined by intracellular and extracellular CAR staining, GFP expression, and confocal microscopy. To understand why the newly designed D.66.α CAR did not express CD4 on the cell surface, our study took a systematic approach to determine the CAR domain(s) affecting surface detection. In doing so, we investigated both the biochemical and spatial requirements of various receptors motifs; an aspect of CAR biology that is still generally poorly understood by the field. Our various CAR modifications identified the CD8α hinge and TM region as the main determinants in CAR expression. Shortening of the hinge from 66 to 45 amino acids partially recovered CAR surface expression, with addition of the LYC motif at the end of the CD8α TM recovering both intracellular and extracellular CD4 expression. In experiments where LYC from the CD8α TM domain was deleted or mutated to TTA or TTC, both extracellular and intracellular CAR expression were severely reduced by flow cytometry and confocal microscopy. Taken together, our results showed that both a shorter hinge and the LYC motif were necessary to have strong CD8αTM CAR surface expression. Furthermore, we determined that CD4-CAR surface expression is maximized by removing the motif FQKAS from the end of the truncated CD4 D1D2 extracellular domain. Finally, we determined though our cytotoxic assay that new CAR vector resulted in CAR T cell killing activity.

The new vectors were designed to reduce the homology of the extracellular CD4 domain while maintaining binding affinity to HIV envelope gp120. The natural CD4 extracellular domain is composed of 4 immunoglobulin (Ig) domains with the N-terminal domain 1 (D1) binding to HIV-1 envelope and soluble D1D2 protein inhibiting viral replication [47]. We eliminated 189 aa from the CD4 (D3D4) and linked the D1D2 Ig domains with a 66- or 45- amino acid extracellular hinge and TM domain from the CD8α receptor. The 66-aa CD8α hinge includes sequences up to the conserved cysteine necessary for the CD8α Ig loop, whereas the 45- aa hinge excludes components of the CD8 Ig domain [46]. We also included 5 aa from D3 (FQKAS) in some vectors as other have previously reported [37]. Since the hinge length and/or sequence may also play a role in processing and stability, presentation of the binding domain, and functional activity, we compared efficacy of the various CAR configurations. Additionally, as others have shown [41], we found that CD4-CARs with CD8α TM domain reduced the susceptibility of CD4-expressing CAR T cell to HIV infection (supplemental data).

Our vector analysis, and the generation of variant CARs, brought to light the requirement for the motif LYC to be included at the end of the CD8α TM. Early mentions of the LYC motif are found in the first studies that attempted to clone CD8α, then known as LYT-2 [48–50]. At the time, since it preceded a stretch of 28 basic amino acids, motif LYC was predicted to be part of the transmembrane domain [51]. Some TM prediction programs also calculate that the LYC will be a part of the TM domain (Supplemental Data). Interestingly, this motif is evolutionarily conserved across species such as human and rat [52]. Additionally, the LYC motif is at the end of CD8α exon 4, which also indicates its evolutionary linkage to the TM domain [46]. However, current crystal structure models predict that the α-helix in the TM precedes LYC and that the LYC motif is part of the CD8α cytoplasmic domain [46]. It remains clear from our experiments that the LYC motif is required for proper receptor surface expression as CARs that lacked this motif had very significantly reduced surface expression. In addition, mutation of the LYC motif resulted in abrogation of surface and intracellular CAR detection. We chose to mutate LYC to TTA and TTC because we aimed to contrast the function of the amino acid residues in LYC. If leucine is a hydrophobic amino acid; then one of its less, but still polar equivalents, is threonine. Furthermore, mutating tyrosine to threonine would remove the aromatic ring and a potential tyrosine phosphorylation site, which could potentially dominate the interaction. Lastly, cysteine was mutated to alanine because their molecular structures are similar while replacing the functional polar CH_2_-SH group with a non-polar CH3. Importantly, the cysteine 206 (C206) in motif LYC of the CD8α gene is necessary for palmitoylation [46], which is involved in the association of membrane proteins and plays an important role in subcellular trafficking [53].

Therefore, in the TTC variant, the cysteine was maintained to determine if its biochemical function was crucial for processing. Since mutating LYC to either TTA or TTC resulted in loss of both extracellular and intracellular CAR expression, it may be that the mutated double threonine affected the proper helix conformation of the upstream transmembrane domain or affected protein trafficking. In support of our data, studies that included only the amino acids LY at the CD8α TM also showed surface expression of a CAR [19], which indicates this motif is important for surface expression. In our experiments, CARs lacking or with a mutated LYC motif had reduced extracellular and intracellular expression, suggesting that its role with CD8α TM is more complex than surface stability. We would predict that modification of the LYC domain on the inside of the TM domain would affect cell membrane localization for most surface antigens, independent of the target, and including scFv-based CAR vectors to tumor antigens.

Aside from the CD8α transmembrane domain, our findings on the length requirements of CD8α hinge for proper CAR expression aligns with that of other studies [19, 54]. The importance of CD8α hinge for receptor surface expression has also been highlighted by other groups; the Williams laboratory synthesized a construct encoding the Ig-like domain of rat CD8α without the CD8α hinge and showed the construct did not express in transfected CHO cells [52]. In our study, CARs with the complete 66 aa CD8α hinge up to the conserved cysteine in the Ig domain (even with the motif LYC present) showed reduced cell surface expression compared to CARs with a shorter 45 aa hinge. In its natural conformation, CD8α has a hinge of 48 aa [52] with 10 aa in the last β strand and 19 aa in the complimentary determining region 3 loop.

Therefore, the longer 66 aa encompassed part of the IgG-like extracellular structural domain of CD8α, which could have affected CAR surface expression due to misfolding. It is important to note that other hinge domains found on anti-HIV CARs are shorter (CD4 hinge (23 aa), CD28 hinge (39 aa), 50 aa [19]) than the natural CD4 receptor, which has a total of 396 aa, or 189 aa in D3D4 linking D1D2 to the TM region. While hinge length may play a role in CAR function, we would also predict hinge domains that incorporate part of the functional domain (ie, an Ig domain) would be less stable and less efficient in surface expression than without it. In a different CAR construct, expression on the cell surface was detected with a 49 aa CD8α hinge [19]. Based on the observation that changes in the TM affected CAR surface expression levels but did not affect CAR mRNA level or total amount of CAR protein, Fujiwara et al. concluded that CD8α hinge affected the transport efficiency of CAR proteins to the cell membrane and that TM regulated the membrane surface stability of CAR [19]. Although both of our studies observed hinge lengths and TM-associate motifs affecting CAR surface expression, our study goes a step further by detecting CAR intracellularly, which allowed us to observe that increasing CD8α hinge length resulted in loss of CAR surface expression but not intracellular expression (Figure 4). Instead, mutation or deletions of motifs affecting the CD8α TM caused loss of both surface and intracellular expression (Figure 4). While more experiments would be necessary to define the mechanisms that affect CAR surface expression, our data suggest that hinge length restriction may affect surface expression stability whereas the transmembrane domain affects protein folding, transport efficiency, or protein degradation.

It is thought that the efficient protein transport through the secretory pathway depends in part on protein folding into a stable structure [55]; and that membrane transport of a CAR is speculated to depend on the folding of its extracellular domain [56]. FQKAS is a motif of 5 aa found at position 204-208 of CD4, which coincides with the beginning of domain 3 but is not part of domain 2 [42, 57]. Therefore, it is possible that the reason we detected less surface expression in functional CARs containing the FQKAS motif at the end of immunoglobulin domain 2 is because its presence negatively affects protein folding (Figure 5 and 6). In addition, as removing FQKAS recovered full expression of the rescued D.45.α^LYC^, it is conceivable that removal of FQKAS from D*.39.28 would have helped recovered expression levels similar to those achieved with the CD8α TM CAR. This finding is of considerable impact for any CAR generated to have only immunoglobulin domain D1 and D2 of CD4 as extracellular domain, especially since the original published CAR sequence included an FQKAS motif [37], which could lower the surface expression of CARs. Even though FQKAS is present in CARs using the full-length CD4 extracellular domain, its presence as a separate entity removed from the rest of a stable structure or a completed domain could affect protein folding and result in lower surface CAR expression.

In either case, expression of CD4 CAR on T cells demonstrated functional activity and were able to kill HIV Env^+^ cells. These *in vitro* assays demonstrate function, but do not necessarily replicate conditions *in vivo*. Other factors, such as the configuration of the intracellular signaling domains, could influence the *in vivo* functional activity such as proliferation or memory cell differentiation. Thus, it may be very difficult to distinguish *in vitro* functional activity of the different various CAR constructs and configurations, or to predict based on the *in vitro* assays which CAR construct will prove efficacious.

At its core, this study underlines the importance of understanding the biochemical structures of various receptor domains as they are evaluated for CAR expression and their subsequent effect on CAR functional activity. Here, like in many CAR structures, the D.66.α CAR was based on the CD4 and CD8 reference sequences; however, these CARs may not account for the possible downstream repercussion of domains and adjacent motifs with various functions as these structures may affect processing and surface expression of different CAR constructs.

In conclusion, by evaluating the mechanisms by which a novel CAR lacked surface expression, this study identified CAR elements in the CD4 extracellular domain, the hinge length, and the CD8α TM region that affect CAR surface expression. These findings not only showcase the importance of understanding how CAR domains and motifs interact within the structure of a receptor, but they also highlight the cellular biochemical machinery that effectively guides successful CAR expression and CAR T cell strategy. Understanding the principles dominating the expression of natural receptors are necessary to generate effective therapies.

Thus, these results contribute to these fundamental guidelines for CAR designs to improve current CAR strategies.

## Material and Methods

### Vector preparations

Vector backbone was shared across all CAR variants used in this study. Briefly, multiple modifications were done to a Moloney Murine Leukemia Virus (MoMLV)-derived vector system including Simian Virus 40 ori/T antigen-mediated episomal replication in packaging cells, replacement of the MoMLV 5’ U3 promoter with a series of stronger composite promoters, and addition of an extra polyadenylation signal downstream of the 3’ long terminal repeat [58]. Variant inserts were synthesized according to the provided amino acid sequences (Supplemental Figure) and cloned by GeneArt (Regensburg, Germany).

### Virus production and transduction

HEK 293T cells were grown in DMEM (Gibco Life Technologies, Grand Island, NY) supplemented with 10% fetal calf serum (Gibco Life Technologies), 1% Penn Strep (Gibco Life Technologies), 2 mM GlutaMax (Gibco Life Technologies), and 25 mM HEPES buffer (Gibco Life Technologies). To generate retroviral particles, HEK 293T cells (1 x10^7^ cells) were plated in D10 media and co-transfected after 18 hours with expression vectors encoding 2 μg VSV glycoprotein, 4 μg MLV Gag/Pol, 4 μg of Rev, and 26 μg the pSRC transfer vectors using calcium phosphate co-precipitation according to manufacturer’s recommendations (Thermofisher Scientific, Waltham, MA). Supernatant was collected from transfected HEK 293T after 48 hours, filtered through 0.45 μm nylon syringe filters, and stored at −80°C.

For transduction, a single cell suspension of HEK 293T cells (0.5×10^6^) were resuspended 2 mL of viral particle supernatant for E:T of 2:1 with 8 μg/mL of Polybrene (Millipore Sigma, Burlington, MA), maintained in at 37°C for 4 hours, and mixed every 20 minutes. Afterwards, cells were plated in a 6-well plate with viral particle supernatant. Media was changed to D10 after 48 hours and cells were cultured for at least another 72 hours before functional assays.

### Vector integration qPCR

Genomic DNA was isolated from transduced HEK 293T cells using the NucleoSpin Tissue Genomic DNA Isolation Kit (Macherey Nagel, Dueren, Germany). For detecting the CAR2 sequences, the forward primer (5’-GCAAGCATTACCAGCCCTAT-3’) and reverse primer (5’- GTTCTGGCCCTGCTGGTA-3’) had a final concentration of 400 nM, while the Probe (5’ 6FAM-ATCGCTCCAGAGTGAAGTTCAGCA-BHQ 3’) had a final concentration of 200 nM, in 25 μL reaction using Taqman Universal Mastermix (Applied Biosystems, Carlsbad, CA) with 100 ng genomic DNA. qPCR was run on Thermofisher 7900HT with the following cycle conditions: 50°C for 2 min, 95°C for 10 min, and 40 cycles of 15 sec at 95°C followed by 64°C for 1 min. To determine copy number, a standard curve was generated consisting of 10^1^ to 10^6^ plasmid copies. Each experimental sample, standard, and NTC was evaluated in triplicate.

### Transfection for functional analysis

To evaluate expression of the CAR vectors, cells transfected directly with the retroviral expression plasmid by calcium-phosphate co-precipitation according to the manufacturer’s protocol (Thermofisher Scientific). Briefly, HEK 293T cells (0.75 x10^6^) were plated in 6-well plates overnight at 37°C and 5%CO_2_. Four hours before transfection, fresh D10 media was exchanged. After 4 hours, the cells were transfected with 10 μg of respective pSCR-CAR plasmid using calcium-phosphate co-precipitation. After 24 hours, an aliquot of cells was taken for confocal microscopy experiments if need be, media was changed, and cells were incubated at 37°C with 5% CO_2_ in incubator for another 24 hours. Cells processed for flow cytometry 48 hours after the initial transfection.

### Flow cytometry

CD4 CAR surface expression was monitored with mouse anti-human CD4-PE, clone RPA-T4 (Biolegends, San Diego, CA), because its epitope is found within the first two domains of CD4. Cells were also stained with a live/dead discriminator dye using the aqua dead cell stain kit (Thermofisher Scientific). Staining protocol was as follows: 2 x10^6^ cells were washed in PBS and stained with live/dead dye according to manufacturer recommendation at RT in the dark for 20 min. For surface staining, cells were then washed with PBS containing 2% fetal bovine serum (FBS) and stained with PE anti-CD4 for 20 min at RT in the dark. Afterwards, cells were washed and fixed with 1% PFA in PBS overnight. For intracellular staining, cells were stained with the live/dead dye, fixed and permeabilized for 20 min with BD Cytofix-Cytoperm and washed according to the manufacturer’s protocol (BD Biosciences, San Jose, CA). Next, cells were stained with PE anti-CD4 for another 20 min at RT in the dark. Cells were washed in PBS and fixed with PBS in 1% PFA overnight. Sample acquisition was performed either on a LSR II (BD Biosciences), LSRFortessa (BD Biosciences), or FACSymphony flow cytometer (BD Biosciences). After gating on live, singlet, a dump channel, and the lymphocyte scatter gate, CD4 detection by PE was assessed. Data was analyzed using FlowJo software (FlowJo LLC, Ashland, OR) and graphed with GraphPad Prism 8 software (GraphPad Software, San Diego, CA).

### Confocal Microscopy

On day 1 post-transfection, 1×10^5^ HEK 293T cells were plated on each respective well of a 4 well chamber slide (Thermofisher Scientific) and incubated overnight at 37°C with 5% CO_2_. The next morning, cells were washed with warm phosphate buffered saline (PBS) and fixed for 30 minutes in 2% paraformaldehyde (PFA) at room temperature. After washing, cells were incubated with 100 mM glycine diluted in PBS +10% normal goat serum (NGS) + 0.02% fish skin gelatin (FSG) + 0.01% triton X100 (TX100) for 20 minutes to block residual PFA. All washes and antibody incubations were done on a rotator platform at room temperature. Cells were washed 3 times in PBS-NGS-FSG-TX100 and incubated for 1 hour with mouse anti-human CD4, clone RPA-T4 (1:100 Invitrogen). Washes were done prior to and following a 1-hour incubation with goat anti-mouse Alexa Fluor 488 (1:1000 Life Technology). Cells were left in wash media overnight at 4°C. The following day, this procedure was repeated using rabbit anti-Caveolin (1:100 Millipore Sigma), and goat anti-rabbit Alexa Fluor 568 (1:1000 Life Technology). Prior to imaging ToPro3 (1:2000 Life Technology), was used to label cellular nuclei. Imaging and image processing was done with a Leica DMi8 (Leica Microsystems, Wetzlar, Germany).

### Retroviral-like particle generation and concentration

HEK 293 T cells were transfected by calcium-phosphate method using vectors pSRC-CAR2 and pSRC-D45a with accessory envelope plasmid pCMV-VSV-G and a packaging plasmid pCMV-Gag/Pol as previously described [58]. Vector particles contained in 10 mL conditioned supernatant from the transfected cells were concentrated through a PEG-mediated precipitation method and resuspended in 100 μl PBMC supplemented with 2% BSA and used to transduce target cells.

### PBMC transduction and expansion

Human peripheral blood mononuclear cells (PBMCs) are isolated from blood samples Gulf Coast Regional Blood Bank, Houston, TX) through ficoll-based density gradient centrifugation, and stimulated and expanded with the T Cell Activation/Expansion Kit (Miltenyi Biotec, Cat No. 130-091-441) according to the vendor’s manual. Three days following the stimulation, 1×10^6^ cells were transduced with the retroviral vectors by resuspending the cell pellet in 100 μl expansion medium and 100 μl concentrated vector. The cell suspension was loaded into Retronectin-coated 24-well plates and incubated at 37°C, 5% CO_2_ for 60 minutes. Subsequently, 1.0 mL cell expansion medium was added to each well and the plate was then centrifuged at 500 xg for 60 minutes at 30°C. After the centrifugation, the transduced PBMC were cultured at 37°C, 5% CO_2_ in humidified incubator, according to the manufacture’s protocol [58].

### Cytotoxic assay

A target HIV envelope positive cell line was stably transduced with a replication-defective lentiviral vector expressing the envelope gene, as well as accessary genes of tat, rev, vpu, and a mRFP reporter gene fused with a puromycin resistant selection marker. Following transduction, the cells were cultured with 2.0 μg/mL puromycin for one week to select for transduced cells.

Cytotoxicity assays were performed in 24 well plate with total cell number of 2.4 x10^6^ cells per well. For the assay with an effector to target ratio of 1:5, 0.4 x10^6^ transduced PBMC were mixed with 2.0 x10^6^ cells of the target cells; for the assay with an effector to target ratio of 1:1, 1.2 x10^6^ transduced PBMC were mixed with 1.2 x10^6^ cells of the target cells; for the assay with an effector to target ratio of 5:1, 2.0 x10^6^ transduced PBMC were mixed with 0.4 x10^6^ cells of the target cells. The cell suspensions were centrifuged at 550 xg for 5 minutes at 4°C, and then resuspended in 1.0 mL T cell expansion medium with 5 μl anti-CD107α-BV785 antibody, inoculated into a 24 well plate and incubated at 37°C cell culture incubator for 24 hours.

Following the 24-hour incubation, the cells were resuspended in 50 μl PBS with 2% BSA and stained with 5 μl anti-CD8-FITC, anti-CD4-PCR-Cy5.5, anti-CD3-APC-H7 antibodies and L/D-BV510 for 30 minutes at 4°C, washed and resuspended using PBS, fixed with 1% paraformadehyde in PBS, and analyzed on the Fortessa Flow Cytometer. Analysis of cytotoxicity was done through analyzing ratio of dead cells among the mRFP positive target cell populations.

### Statistics

*In vitro* CAR expression significance was detected using an unpaired two-sample *t* test comparing 2 groups at a time (CAR vs. CAR extracellular or intracellular). Significant results are reported on each figure (p values: >0.05, *≤0.05, **≤0.01, ***≤0.0001). Analysis was performed on GraphPad Prism 8 software.

## Acknowledgment

We thank Cariappa Annaiah for his suggestions and mentorship.

## Author contribution

G.Z. and S.E.B. conceived the project, designed experiments, interpreted results, and wrote the manuscript. A.K. designed experiments and reviewed and interpreted results. C.W. generated the D.66.α CAR construct and performed the primary T cell transductions and the cytotoxic assay. C.C.M. and C.P.G. performed confocal/fluorescent microscopy experiments. N.M.J. conducted qPCR experiments and analysis. W.W. interpreted the results.

## References

1. Wallstabe, L., et al., ROR1-CAR T cells are effective against lung and breast cancer in advanced microphysiologic 3D tumor models. JCI Insight, 2019. 4(18).

2. Zhen, A., et al., Long-term persistence and function of hematopoietic stem cell-derived chimeric antigen receptor T cells in a nonhuman primate model of HIV/AIDS. PLoS Pathog, 2017. 13(12): p. e1006753.

3. Benmebarek, M.R., et al., Killing Mechanisms of Chimeric Antigen Receptor (CAR) T Cells. Int J Mol Sci, 2019. 20(6).

4. Porcellini, S., et al., CAR T Cells Redirected to CD44v6 Control Tumor Growth in Lung and Ovary Adenocarcinoma Bearing Mice. Front Immunol, 2020. 11: p. 99.

5. Brudno, J.N., et al., Safety and feasibility of anti-CD19 CAR T cells with fully human binding domains in patients with B-cell lymphoma. Nat Med, 2020. 26(2): p. 270–280.

6. Srivastava, S. and S.R. Riddell, Chimeric Antigen Receptor T Cell Therapy: Challenges to Bench-to-Bedside Efficacy. J Immunol, 2018. 200(2): p. 459–468.

7. Mohanty, R., et al., CAR T cell therapy: A new era for cancer treatment (Review). Oncol Rep, 2019. 42(6): p. 2183–2195.

8. Abramson, J.S., Anti-CD19 CAR T-Cell Therapy for B-Cell Non-Hodgkin Lymphoma. Transfus Med Rev, 2020. 34(1): p. 29–33.

9. Kallam, A. and J.M. Vose, Recent Advances in CAR-T Cell Therapy for Non-Hodgkin Lymphoma. Clin Lymphoma Myeloma Leuk, 2019. 19(12): p. 751–757.

10. Chavez, J.C., C. Bachmeier, and M.A. Kharfan-Dabaja, CAR T-cell therapy for B-cell lymphomas: clinical trial results of available products. Ther Adv Hematol, 2019. 10: p. 2040620719841581.

11. Brudno, J.N. and J.N. Kochenderfer, Recent advances in CAR T-cell toxicity: Mechanisms, manifestations and management. Blood Rev, 2019. 34: p. 45–55.

12. Liu, E., et al., Cord blood NK cells engineered to express IL-15 and a CD19-targeted CAR show long-term persistence and potent antitumor activity. Leukemia, 2018. 32(2): p. 520–531.

13. Shah, N.N. and T.J. Fry, Mechanisms of resistance to CAR T cell therapy. Nat Rev Clin Oncol, 2019. 16(6): p. 372–385.

14. Zenere, G., et al., Optimizing intracellular signaling domains for CAR NK cells in HIV immunotherapy: a comprehensive review. Drug Discov Today, 2019. 24(4): p. 983–991.

15. Ma, S., et al., Current Progress in CAR-T Cell Therapy for Solid Tumors. Int J Biol Sci, 2019. 15(12): p. 2548–2560.

16. Scholler, J., et al., Decade-long safety and function of retroviral-modified chimeric antigen receptor T cells. Sci Transl Med, 2012. 4(132): p. 132ra53.

17. Abate-Daga, D. and M.L. Davila, CAR models: next-generation CAR modifications for enhanced T-cell function. Mol Ther Oncolytics, 2016. 3: p. 16014.

18. Navai, S.A. and N. Ahmed, Targeting the tumour profile using broad spectrum chimaeric antigen receptor T-cells. Biochem Soc Trans, 2016. 44(2): p. 391–6.

19. Fujiwara, K., et al., Structure of the Signal Transduction Domain in Second-Generation CAR Regulates the Input Efficiency of CAR Signals. Int J Mol Sci, 2021. 22(5).

20. Alabanza, L., et al., Function of Novel Anti-CD19 Chimeric Antigen Receptors with Human Variable Regions Is Affected by Hinge and Transmembrane Domains. Mol Ther, 2017. 25(11): p. 2452–2465.

21. Zhong, X.S., et al., Chimeric antigen receptors combining 4-1BB and CD28 signaling domains augment PI3kinase/AKT/Bcl-XL activation and CD8+ T cell-mediated tumor eradication. Mol Ther, 2010. 18(2): p. 413–20.

22. Yeku, O.O., et al., Armored CAR T cells enhance antitumor efficacy and overcome the tumor microenvironment. Sci Rep, 2017. 7(1): p. 10541.

23. Scarfo, I. and M.V. Maus, Current approaches to increase CAR T cell potency in solid tumors: targeting the tumor microenvironment. J Immunother Cancer, 2017. 5: p. 28.

24. Heczey, A., et al., CAR T Cells Administered in Combination with Lymphodepletion and PD- 1 Inhibition to Patients with Neuroblastoma. Mol Ther, 2017. 25(9): p. 2214–2224.

25. Yang, O.O., et al., Lysis of HIV-1-infected cells and inhibition of viral replication by universal receptor T cells. Proc Natl Acad Sci U S A, 1997. 94(21): p. 11478–83.

26. Sahu, G.K., et al., Anti-HIV designer T cells progressively eradicate a latently infected cell line by sequentially inducing HIV reactivation then killing the newly gp120-positive cells. Virology, 2013. 446(1-2): p. 268–75.

27. Walker, R.E., et al., Long-term in vivo survival of receptor-modified syngeneic T cells in patients with human immunodeficiency virus infection. Blood, 2000. 96(2):: p. 467–74.

28. Mitsuyasu, R.T., et al., Prolonged survival and tissue trafficking following adoptive transfer of CD4zeta gene-modified autologous CD4(+) and CD8(+) T cells in human immunodeficiency virus-infected subjects. Blood, 2000. 96(3): p. 785–93.

29. Deeks, S.G., et al., A phase II randomized study of HIV-specific T-cell gene therapy in subjects with undetectable plasma viremia on combination antiretroviral therapy. Mol Ther, 2002. 5(6): p. 788–97.

30. Finney, H. and e. al, Chimeric receptors providing both primary and costimulatory signaling in T cells from a single gene product. J Immunol, 1998. 161(6):: p. 2791–7.

31. Kawalekar, O.U., et al., Distinct Signaling of Coreceptors Regulates Specific Metabolism Pathways and Impacts Memory Development in CAR T Cells. Immunity, 2016. 44(2): p. 380–90.

32. Campana, D., H. Schwarz, and C. Imai, 4-1BB chimeric antigen receptors. Cancer J, 2014. 20(2): p. 134–40.

33. Kwong, P.D., et al., Structure of an HIV gp120 envelope glycoprotein in complex with the CD4 receptor and a neutralizing human antibody. Nature, 1998. 393(6686): p. 648–59.

34. Traunecker, A., W. Luke, and K. Karjalainen, Soluble CD4 molecules neutralize human immunodeficiency virus type 1. Nature, 1988. 331(6151): p. 84–6.

35. Ryu, S.E., et al., Crystal structure of an HIV-binding recombinant fragment of human CD4. Nature, 1990. 348(6300): p. 419–26.

36. Sharma, D., et al., Protein minimization of the gp120 binding region of human CD4. Biochemistry, 2005. 44(49): p. 16192–202.

37. Liu, L., et al., Novel CD4-Based Bispecific Chimeric Antigen Receptor Designed for Enhanced Anti-HIV Potency and Absence of HIV Entry Receptor Activity. J Virol, 2015. 89(13): p. 6685–94.

38. Sakihama, T., A. Smolyar, and E.L. Reinherz, Oligomerization of CD4 is required for stable binding to class II major histocompatibility complex proteins but not for interaction with human immunodeficiency virus gp120. Proc Natl Acad Sci U S A, 1995. 92(14): p. 6444–8.

39. Moebius, U., et al., Delineation of an extended surface contact area on human CD4 involved in class II major histocompatibility complex binding. Proc Natl Acad Sci U S A, 1993. 90(17): p. 8259–63.

40. Cerutti, N., et al., Disulfide reduction in CD4 domain 1 or 2 is essential for interaction with HIV glycoprotein 120 (gp120), which impairs thioredoxin-driven CD4 dimerization. J Biol Chem, 2014. 289(15): p. 10455–10465.

41. Leibman, R.S., et al., Supraphysiologic control over HIV-1 replication mediated by CD8 T cells expressing a re-engineered CD4-based chimeric antigen receptor. PLoS Pathog, 2017. 13(10): p. e1006613.

42. CD4 molecule [human]. https://www.ncbi.nlm.nih.gov/gene/920. National Center for Biotechnology Information, National Library of Medicine. 10-Oct-2023.

43. Maclean, A., et al., A Novel Real-Time CTL Assay to Measure Designer T Cell Function Against HIV Env+ Cells. J Med Primatology, 2014. 43(5): p. 341–8.

44. Liu, Z., et al., Systematic comparison of 2A peptides for cloning multi-genes in a polycistronic vector. Sci Rep, 2017. 7(1): p. 2193.

45. Lagenaur, L.A., et al., sCD4-17b bifunctional protein: extremely broad and potent neutralization of HIV-1 Env pseudotyped viruses from genetically diverse primary isolates. Retrovirology, 2010. 7: p. 11.

46. CD8 subunit alpha [human]. https://www.ncbi.nlm.nih.gov/gene/925. National Center for Biotechnology Information, National Library of Medicine. 15-Oct-2023.

47. Lu, L., et al., A bivalent recombinant protein inactivates HIV-1 by targeting the gp41 prehairpin fusion intermediate induced by CD4 D1D2 domains. Retrovirology, 2012. 9: p. 104.

48. Blobel, G., et al., Translocation of proteins across membranes: the signal hypothesis and beyond. Symp Soc Exp Biol, 1979. 33: p. 9–36.

49. Walker, I.D., et al., Ly antigens associated with T cell recognition and effector function. Immunol Rev, 1984. 82: p. 47–77.

50. Luescher, B., et al., The mouse Lyt-2/3 antigen complex--I. Mode of association of the subunits with the membrane. Mol Immunol, 1984. 21(4): p. 329–36.

51. Zamoyska, R., et al., Two Lyt-2 polypeptides arise from a single gene by alternative splicing patterns of mRNA. Cell, 1985. 43(1): p. 153–63.

52. Classon, B.J., et al., The hinge region of the CD8 alpha chain: structure, antigenicity, and utility in expression of immunoglobulin superfamily domains. Int Immunol, 1992. 4(2): p. 215–25.

53. Pang, D.J., A.C. Hayday, and M.J. Bijlmakers, CD8 Raft localization is induced by its assembly into CD8alpha beta heterodimers, Not CD8alpha alpha homodimers. J Biol Chem, 2007. 282(18): p. 13884–94.

54. Guedan, S., et al., Engineering and Design of Chimeric Antigen Receptors. Mol Ther Methods Clin Dev, 2019. 12: p. 145–156.

55. Lippincott-Schwartz, J., T.H. Roberts, and K. Hirschberg, Secretory protein trafficking and organelle dynamics in living cells. Annu Rev Cell Dev Biol, 2000. 16: p. 557–89.

56. Krug, C., et al., Stability and activity of MCSP-specific chimeric antigen receptors (CARs) depend on the scFv antigen-binding domain and the protein backbone. Cancer Immunol Immunother, 2015. 64(12): p. 1623–35.

57. Lange, G., et al., Crystal structure of an extracellular fragment of the rat CD4 receptor containing domains 3 and 4. Structure, 1994. 2(6): p. 469–81.

58. Wu, C. and Y. Lu, High-titre retroviral vector system for efficient gene delivery into human and mouse cells of haematopoietic and lymphocytic lineages. J Gen Virol, 2010. 91(Pt 8): p. 1909–1918.

